# Muscle-restricted Nox4 knockout partially corrects muscle contractility following spinal cord injury in mice

**DOI:** 10.1101/2023.08.04.551985

**Authors:** Carlos A. Toro, Rita De Gasperi, Abdurrahman Aslan, Nicholas Johnson, Mustafa M. Siddiq, Christine Chow, Wei Zhao, Lauren Harlow, Zachary Graham, Xin-Hua Liu, Junichi Sadoshima, Ravi Iyengar, Christopher P Cardozo

## Abstract

Spinal cord injury (SCI) results in severe atrophy of skeletal muscle in paralyzed regions, and a decrease in the force generated by muscle per unit of cross-sectional area. Oxidation of skeletal muscle ryanodine 1 receptors (RyR1) reduces contractile force due to reduced binding of calstabin 1 to RyR1 together with altered gating of RyR1. One cause of RyR1 oxidation is NADPH oxidase 4 (Nox4). We have previously shown that in rats, RyR1 was oxidized and bound less calstabin 1 at 56 days after spinal cord injury (SCI) by transection. Here, we used a conditional knock-out mouse model of Nox4 in muscle to investigate the role of Nox4 in reduced muscle specific force after SCI. Peak twitch force in control mice after SCI was reduced by 42% compared to sham-operated controls but was increased by approximately 43% in SCI Nox4 conditional KO mice compared to SCI controls although it remained less than that for sham-operated controls. Unlike what observed in rats, after SCI the expression of Nox4 was not increased in gastrocnemius muscle and binding of calstabin 1 to RyR1 was not reduced in this muscle. The results suggest a link between Nox4 expression in muscle tissue and reduction in muscle twitch force, however further studies are needed to understand the mechanistic basis for this linkage.

## INTRODUCTION

Spinal cord injury (SCI) results in sub-lesional paralysis and atrophy of skeletal muscles. In rodents, 40 to 60% of muscle mass is lost within the first few weeks after SCI [1-3]. A similar though slower decrease has been reported in humans based on analysis of declines in muscle fiber cross sectional area over the first six months after SCI [4]. Muscle paralyzed by SCI is weaker and more easily fatigued. Rat soleus muscle was found to have a reduction in absolute force generation at 3 and 6 months after spinal cord transection associated with faster fatigue, a slow to fast transition of myosin heavy chain isoforms and faster rates of contraction [5]. Single fiber studies of rat tibialis anterior (TA) muscle at 2 and 4 weeks after a contusion SCI revealed reduced absolute force at 2 weeks post-SCI and reduced specific force (force produced per unit area) at 2 and 4 weeks after injury [6].

The reasons for this reduced specific force generation after SCI are poorly understood. One possibility is impaired function of ryanodine receptor 1 (RyR1) in skeletal muscle. RyR1 is a large calcium activated calcium channel present in the membrane of the endoplasmic/sarcoplasmic reticulum [7-9]. In skeletal muscle, RyR1 is localized to the triad, a cellular structure formed by the junction of evaginations of the sarcoplasmic reticulum with T-tubules through formation of complexes of dihydropyridine receptors (L-type voltage-gated calcium channels localized to the cytoplasmic membrane/sarcolemma) and RyR1 [7-9]. Depolarization of the cytoplasmic membrane results in opening of RyR1 and release of calcium stored in the sarcoplasmic reticulum resulting in muscle contraction in a process referred to as excitation contraction coupling (ECC).

The open probability of RyR1 is regulated by several factors including binding of calstabin 1, a small protein that stabilizes RyR1 in a closed state [10]. Oxidation and nitrosylation of RyR1 decrease the binding of calstabin 1 to RyR1 and are associated with frequent bursts of calcium release through individual RyR1 channels (i.e., calcium sparks) and are thought to deplete sarcoplasmic reticulum calcium stores to reduce effectiveness of muscle contraction in response to neural activation of skeletal muscle. Administration of molecules that restore normal binding of calstabin 1 to RyR1 significantly improves muscle contractile function in animal models in which there is extensive oxidation and nitrosylation of skeletal muscle RyR1 [11-13]. The source of the reactive oxygen species (ROS) responsible for oxidative modifications of RyR1 may include NADPH+ oxidases (Nox). Nox are widely expressed enzymes thought to be present in virtually all eukaryotic cells [14-17]. They oxidize NADPH+ to generate ROS in response to physiological cues and participate in many physiological processes such as insulin action, regulation of vascular tone, ECC and many others [15-17]. Six Nox isoforms are known [14]. Nox4 is present in skeletal muscle where it has been found to form complexes with RyR1, and participates in oxidation in proportion to oxygen levels thereby potentiating RyR1 opening and muscle contraction as oxygen becomes more abundant [18]. Excessive expression of Nox4 mediates RyR1 oxidation and muscle weakness in mouse models of breast cancer bone metastases [19]. While Nox4 was initially believed to be constitutively active, more recent literature indicates that its activity is tightly regulated [16] by tissue oxygen levels, cellular NADPH levels, and regulatory factors that include interactions with polymerase delta-interacting protein 2 (Poldip2) [20].

The possibility that muscle RyR1 becomes oxidized or nitrosylated following SCI has been addressed in a rat model of spinal cord transection Our group has previously demonstrated immunoprecipitated RyR1 isolated from rat gastrocnemius muscle at 56 days after SCI had elevated levels of oxidation and nitrosylation associated with almost complete loss of calstabin 1 binding [21]. Expression of Nox4 was increased in gastrocnemius muscle at 56 days after SCI and amounts of Nox4 co-immunoprecipitated with RyR1 were elevated after SCI [21], suggesting a mechanistic linkage between upregulation of Nox4 expression and RyR1 oxidation and nitrosylation. In the present study, we tested the hypothesis that upregulation of Nox4 is responsible for reduced specific force of skeletal muscle paralyzed by SCI. We conditionally knockedout Nox4 in cells of the myogenic lineage expressing the myogenic differentiation factor MyoD [22, 23], using a mouse line in which the coding sequence for MyoD was replaced with that for Cre thereby driving Cre expression via the full length murine MyoD promoter [24].

## METHODS

### Animals

Animals were housed in a temperature-controlled vivarium at approximately 20 °C degrees with 12:12 hour light-dark cycles and were provided standard mouse chow and water ad libitum. All animal studies were conducted in conformance with the requirements of The Guide for Care and Use of Laboratory Animals and all other applicable regulations. Animal studies were approved by the IACUC at the James J Peters VA Medical Center (Protocol CAR-17-23), a AAALAC (Association for Assessment and Accreditation of Laboratory Animal Care) accredited facility.

### Development of Mice with Conditional Inactivation of Nox4 in Skeletal Muscle (cKO)

A commercially available line in which the coding sequence for one allele of MyoD was replaced with that for Cre (FVB.Cg-Myod1tm2.1(icre)Glh/J; Jackson Laboratories Stock No: 014140; MyoD-Cre) [24] was used to knockout Nox4 in cells of the myogenic lineage, including mature, adult muscle fibers. Mice heterozygous for MyoD-Cre were crossed with mice in which exon 9 of the Nox4 gene is flanked by lox P sites (Nox4(f/f)) [25] which was generously provided by Dr. Junichi Sadoshima (Rutgers New Jersey Medical School). Excision of exon 9 results in a truncated Nox4 protein that lacks catalytic activity.

### Genotyping and Confirmation of Recombination

Genotyping was performed using genomic DNA isolated from ear snips obtained at the time of weaning. For detection of the Nox4 floxed allele, the following primers were used: primer 102: 5’ GCACTATGCCGAATTGCTCT, primer 103: 5’GAATGCACCGAGCACATTTG and primer 105: 5’ GAGGCTATTCGGCTATGACT. The wild-type allele and the floxed allele were genotyped in separate PCR reactions combining primer 102 and 103 for the wild-type allele to generate a 900 bp band and primers 102 and 105 for the floxed allele to generate a 1300 bp band. The reaction was performed with TopTaq polymerase and buffer (Qiagen) in the presence of 2 mM MgCl_2_ under the following conditions: 95°C for 3 min; 32 cycles at 94°C for 30 sec, 55°C for 30 sec, 72°C for 1 min 30 sec; 72°C for 5 minutes. Genotyping for presence of the MyoD Cre allele was performed using the following primers: MyoD-Cre -wt-F: 5’ CGGCTACCCAAGGTGGAGAT; MyoD-Cre-Mut-F: 5’ GCGGATCCGAATTCGAAGTTCC; Myo-Cre-R: 5’ TGGGTCTCCAAAGCGACTCC. The reaction was performed with TopTaq polymerase and buffer (Qiagen) in the presence of 2 mM MgCl_2_ under the following conditions: 95°C for 3 min; 35 cycles at 94°C for 30 sec, 60°C for 45 sec, 72°C for 45 sec; 72°C for 10 minutes. The PCR products were 343 bp for wild type and 149 bp for the mutant allele; both bands were present for heterozygous mice.

### Experimental Design

Nox4 cKO mice or littermates carrying the floxed Nox4 (Nox4(f/f); genotype controls) gene at 4 to 6 months of age were used. Mice were randomly assigned to either spinal cord transection or sham transection surgeries. At 56 days after surgery, animals were deeply anesthetized by inhalation of isoflurane and the left extensor digitorum longus (EDL) was removed by careful dissection then subjected to ex-vivo physiological testing. The remaining muscles were carefully removed by dissection, weighed and snap frozen in liquid nitrogen.

### SCI Surgeries and Post-operative Care

Animals were randomly assigned to Sham or SCI surgeries, weighed and anesthetized by inhalation of isoflurane. Hair along the spine was removed with a clipper and skin was cleaned with 70% ethanol and beta-iodine solution. After exposing the spine through a midline incision centered at the T9-T10 intervertebral space, vertebral arches at T9-T10 were exposed by careful dissection and the T9 vertebral arch was carefully removed with sharp scissors. For animals assigned to the SCI group, a 0.1 ml of lidocaine solution were applied to the dura to anesthetize the area of injury, after which the spinal cord was severed with sharp microscissors. Completeness of transection was verified visually by retraction of ends of the severed cord. A small piece of surgical sponge was placed between the ends of the cord. Wounds were closed in layers using suture. Animals were administered carprofen and Baytril preemptively after anesthesia but before the incision then for 3 days thereafter. Urine was expressed manually 2 times daily until spontaneous void of the bladder was observed. Thereafter, bladders were checked daily.

### Tissue Harvest and Ex-vivo Physiology

Animals were weighed then anesthetized using inhaled isoflurane. Hindlimb muscles were excised after careful blunt dissection. Measurement of whole-muscle contractile and mechanical properties was performed using an Aurora Scientific ex-vivo physiology system for mice (Aurora, Ontario, Canada). Contractile properties of the EDL muscle was evaluated. Briefly, a 4-0 silk suture was tied to the proximal and distal tendons of intact right EDL, immediately distal to the aponeuroses. Following suture placement, muscles were then removed from the animal and immediately placed in a bath containing a Krebs mammalian Ringer solution at pH 7.4, supplemented with tubocurarine chloride (0.03 mM) and glucose (11 mM) for 10 min. The bath was maintained at 25°C and bubbled constantly with a mixture of O2 (95%) and CO2 (5%). The distal tendon of the muscle was then tied to a dual-mode servomotor/force transducer (Aurora Scientific, Aurora, Ontario, Canada) and the proximal tendon tied to a fixed hook. Using wave pulses delivered from platinum electrodes connected to a high-power bi-phasic current stimulator (Aurora Scientific, Aurora, Ontario, Canada) each EDL was stimulated to contract. The 610A Dynamic Muscle Control v5.5 software (Aurora Scientific, Aurora, Ontario, Canada) was used to control pulse properties and servomotor activity, and record data from the force transducer. Optimal voltage was determined by increasing the stimulation voltage of a single twitch pulse until force did not increase. This voltage was then doubled to ensure maximal stimulus. Optimal length (Lo) was then established by lengthening each EDL by 0.5 mm until force no longer increase following step-wise lengthening. Muscle was allowed to rest for 45 s between twitch responses. Following determination of Lo, muscle was allowed to rest for 45 s and we collected twitch force. After a two-minute rest, we then established the frequency-force relationship. Here, EDL were stimulated at increasing frequencies (i.e. 10, 25, 40, 60, 80, 100, and 150 Hz). Stimulation was delivered for 300 ms, and muscles were left to rest for 1 min between successive stimuli. Maximum absolute isometric tetanic force (Po) was determined from the plateau of the frequency-force relationship. Muscles were then removed from the bath solution and weighed. All data collected were analyzed using the Dynamic Muscle Analysis v5.3 software (Aurora Scientific, Aurora, Ontario, Canada).

### Immunoprecipitation and Western Blotting

Immunoprecipitation of RyR1 and Western blotting were performed as previously described [11]. Gastrocnemius muscle was homogenized in 1 ml of 50 mM Tris HCl pH 7.4, 015M NaCl, 0.5% Triton X-100 supplemented with Halt protease and phosphatase inhibitors (ThermoFisher). The homogenate was centrifuged at 14,000 rpm for 20 minutes and the supernatant saved. Protein concentrations were determined using the BCA reagent (ThermoFisher). For immunoprecipitation (IP), 0.5 mg protein was precleared for 30 minutes at 4°C with 4% Agarose beads (ThermoFisher). To immunoprecipitate RYR1, 1.2 μg of anti-RYR1 monoclonal antibody (ThermoFisher MA 3-925,) was added to the precleared extracts and incubated 1h at 4C. The samples were incubated with Protein A/G-agarose (Santa Cruz) for 1 h at 4°C. The beads were washed three times with the lysis buffer and heated at 95°C for 5 minutes. The immunoprecipitated proteins were separated by SDS-PAGE and transferred to a PVDF membrane. After incubation in blocking buffer (Tris-buffered saline 0.5%, 0.1%Tween-20 (TBST), 0.5 % non-fat dry milk the membrane was incubated overnight with rabbit polyclonal anti-calstabin 1 (PA 1-026, ThermoFisher) (1:1200 in blocking buffer). The membrane was washed with TBST and incubated for 1.5 h with HRP-conjugated anti rabbit IgG (1:10,000 in blocking buffer (Cytiva) and the bands revelated with ECL Prime reagent (Cytiva). To determine the amount of immunoprecipitated RYR, the membrane was stripped and reprobed with the mouse monoclonal anti-RYR1 receptor antibody (MA 3-925, 1:2500). For Nox4 analysis, proteins (50 μg) were separated by SDS Page and blotted onto PVDF membranes. The membranes were probed with rabbit polyclonal anti Nox4 (sc-30141, Santa Cruz) (1:500 dilution) and reprobed with a mouse monoclonal anti-GAPDH (GT239, GeneTex) (1:4000) as loading control. Blots were imaged with an ImageQuant 800 (Cytiva) and quantitated using the Image Quant TL software package (Cytiva).

### Enzyme-linked Immunosorbent Assay (ELISA) to Detect Protein Carbonyl Residues

Tibialis anterior (TA) muscle was homogenized with ceramic spheres in the Fastprep-24 tissue homogenizer (MP Biomedicals, OH, USA) using 20 μl/mg RIPA buffer supplemented with a protease/phosphatase inhibitor cocktail (Cell Signaling Technology, MA, USA). The homogenized tissue was centrifuged at 14,000rpm at 4°C for 10 minutes, and the supernatant was collected for analysis. Total protein concentration was calculated with the Bradford Assay (BioRad). The tissue lysates were treated with 1% DNase (Qiagen, Hilden, Germany) and diluted to a protein concentration of 10μg/ml. Protein oxidation levels were quantified using an OxiSelect Protein Carbonyl ELISA Kit (Cell Biolabs, CA, USA) according to the manufacturer’s instructions.

### Data and Statistical Analysis

Statistical calculations were performed using Graphpad Prism 9. Data are expressed as mean values ± SEM and were analyzed by 2-way mix model ANOVA (knockout x SCI) with a Tukey’s test post-hoc to test for differences between pairs of means. A 2-way ANOVA and multiple comparison analysis were performed using Matlab software (Mathworks, MA, USA) for ELISA.

## RESULTS

Animal body weights determined at both the day of surgery and the time of euthanasia (Fig. 1) were similar between SCI Nox4 cKO and SCI Nox4(f/f) littermates. Body weights were reduced in the SCI groups regardless of genotype when compared to control sham operated mice at time of euthanasia (Fig. 1 B, C). Nox4 cKO and Nox4(f/f) mice seemed to experience a similar degree of body weight loss after SCI (Fig. 1C).

**Figure 1.**
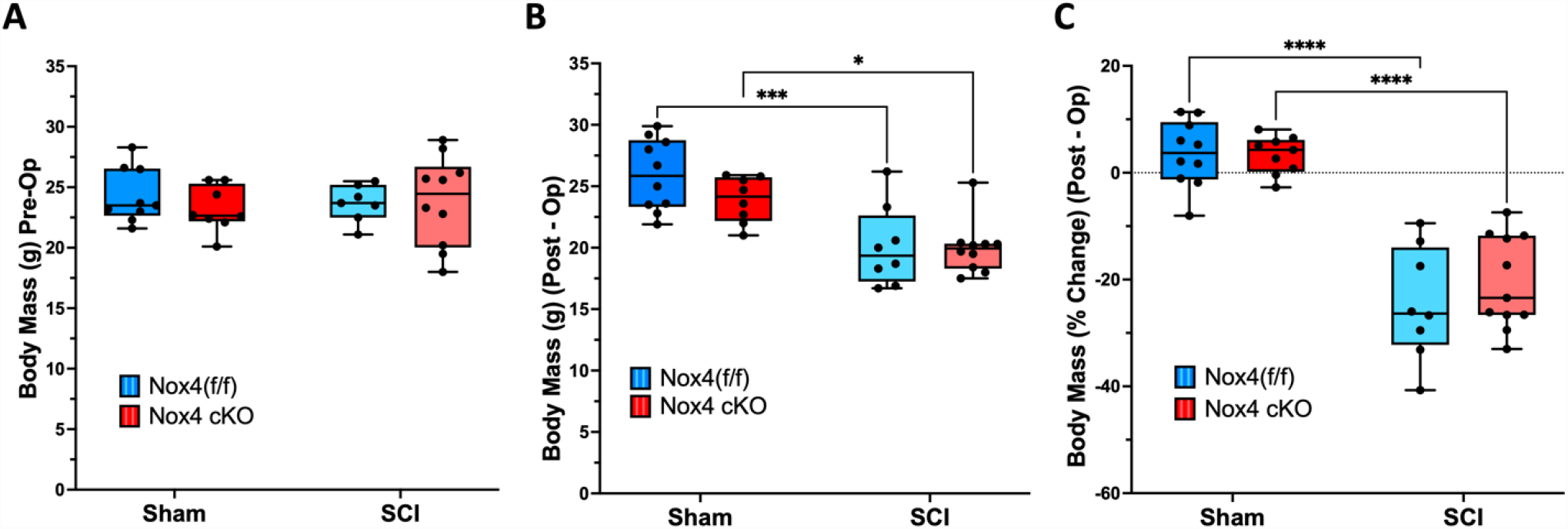
Changes in body mass after SCI. Body weights were determined at A) the day of surgery (Pre-operatively) and B) at time of euthanasia (post-operatively). In C), percent changes in body weight were compared to pre-operative body weight. Dark and light blue bars indicate Nox4 (f/f) Sham and SCI, respectively. Red and pink bars indicate Nox4 cKO Sham and SCI, respectively. Data were analyzed by mixed-model 2-way ANOVA with a Sidak’s multiple comparisons test post-hoc. **, P < 0.01; ***, P < 0.001; ****, P < 0.0001.

Muscle weights were determined at 56 days after SCI and normalized to pre-operative body weight. For gastrocnemius muscle, normalized weights were similar between sham-operated Nox4 cKO and Nox4(f/f) littermate controls (Fig. 2A). Normalized gastrocnemius weights were significantly reduced in SCI Nox4 cKO and SCI Nox4(f/f) groups compared to sham operated controls (Fig. 2A). There was no difference in muscle weight between SCI Nox4 cKO and SCI Nox4(f/f) littermate controls (Fig. 2A). The same pattern was observed for other lower hindlimb muscles including TA, EDL, soleus and plantaris (Fig. 2B-E). Weights of triceps muscles, a weight-bearing forelimb muscle that was not paralyzed by the SCI, were also determined. Triceps weights were similar between sham Nox4 cKO and sham Nox4(f/f) mice, and between SCI Nox4 cKO and SCI Nox4(f/f) mice (Fig. 2F). Triceps weights were significantly reduced in SCI Nox4 cKO and SCI Nox4(f/f) mice compared to sham operated controls (Fig. 2F). In summary, Nox4 cKO did not alter muscle mass in sham-operated mice or alter muscle atrophy in SCI mice at the 56-day timepoint studied.

**Figure 2.**
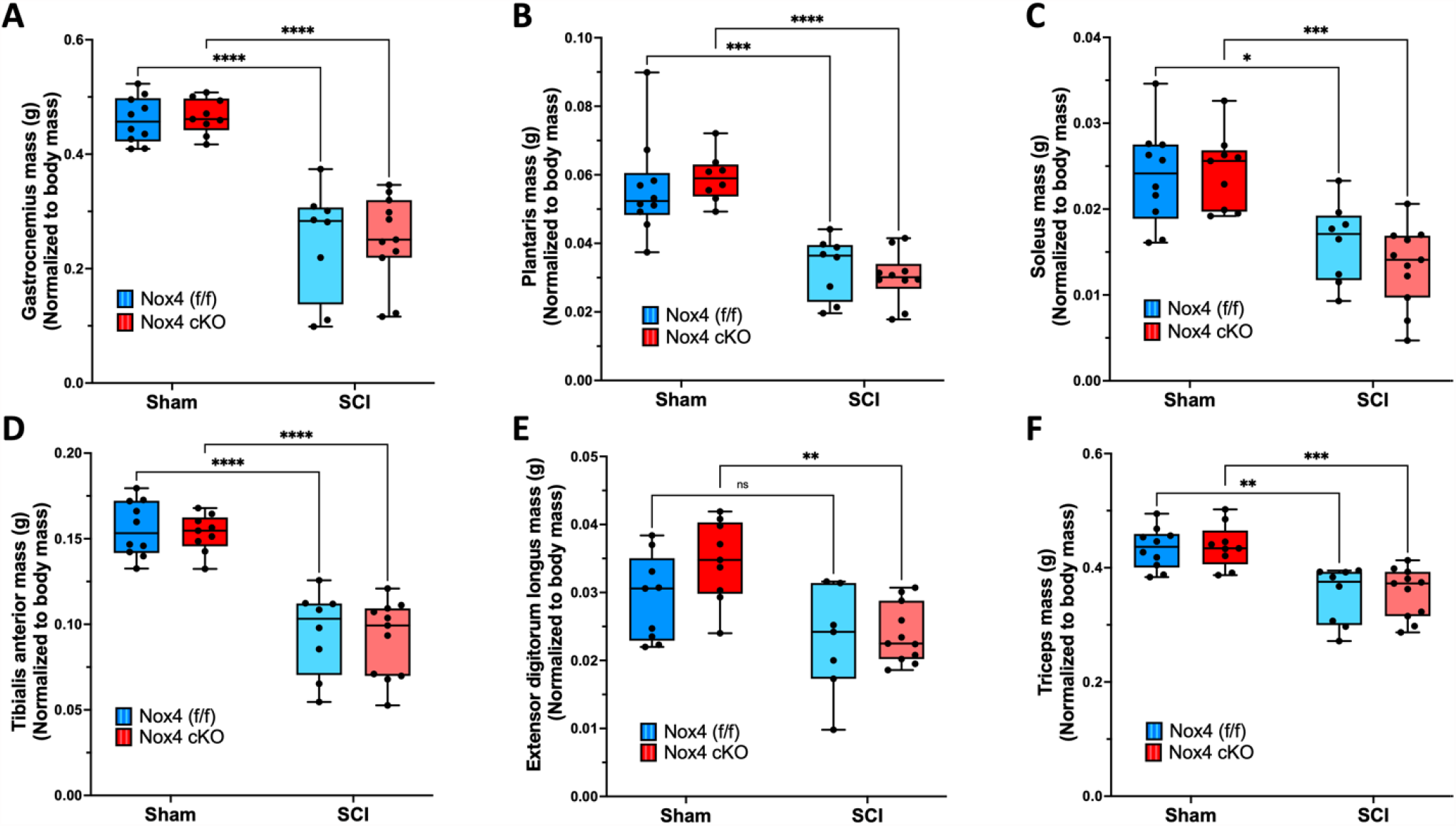
Weights of skeletal muscles are shown after normalization relative to pre-operative body weight. Normalized weights of A) Gastrocnemius; B) Plantaris; C) Soleus; D) Tibialis anterior; E) Extensor digitorum longus; and F) Triceps muslces are shown. Dark and light blue bars indicate Nox4 (f/f) Sham and SCI, respectively. Red and pink bars indicate Nox4 cKO Sham and SCI, respectively. Data were analyzed by mixed-model 2-way ANOVA with a Sidak’s multiple comparisons test post-hoc. *, P < 0.05; **, P < 0.01; ***, P < 0.001.

To determine whether the Nox4 cKO improved muscle force generating capacity after SCI, EDL muscles were used for ex-vivo physiological studies. Analysis of force-frequency curves revealed a significant main-effect for genotype in the SCI groups (P < 0.0001). Force generated by SCI mice with the Nox4 cKO was significantly increased as compared to SCI genotype controls (Nox4(f/f)) at 60, 80, 100 and 150 Hz (Fig. 3A). Peak twitch tension tended to be lower in sham Nox4 cKO versus sham Nox4(f/f) mice although this difference did not reach the threshold for significance (Fig. 3B). Peak twitch tension was significantly reduced for SCI Nox4(f/f) compared to sham Nox4(f/f) controls (Fig. 3B). Peak twitch tension was higher for SCI Nox4 cKO compared to SCI Nox4(f/f) and tended to be lower than sham operated controls although this difference was not significant (Fig. 3B). Time to peak twitch tension was not different between sham Nox4 cKO and sham Nox4(f/f) groups and was not altered by SCI (Fig. 3C). Half-relaxation time was also not different between any of the groups (Fig. 3D). An unexpected pattern in the data was reduced muscle force generation in the sham Nox4 cKO mice compared to sham Nox4(f/f) mice when EDL muscle was tested by force frequency studies or analysis of twitch contractions (Fig. 3A and 3B); these differences did not reach the threshold for significance. In summary, the Nox4 cKO improved EDL force generation during twitch and did not appreciably alter time to peak tension or half-relaxation time.

**Figure 3.**
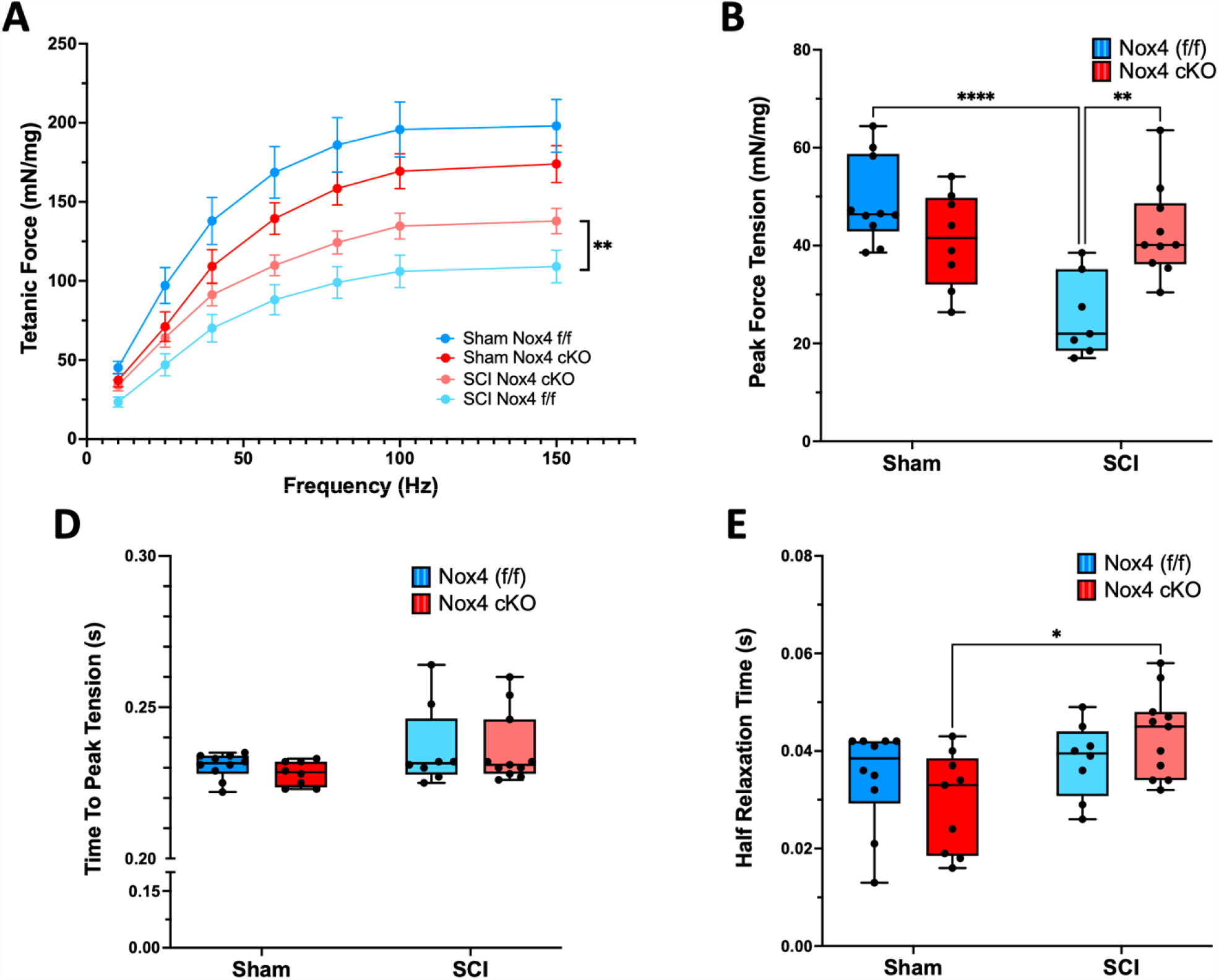
Ex-vivo physiological properties of EDL muscles are shown: A) Tetanic force-frequency was analyzed; B) Peak tetanic tension was measured at 150 Hz; C) Time to peak tension was determined; and D) Half relaxation time was assessed. Dark and light blue bars indicate Nox4 (f/f) Sham and SCI, respectively. Red and pink bars indicate Nox4 cKO Sham and SCI, respectively. Data were analyzed by mixed-model 2-way ANOVA with a Sidak’s multiple comparisons test post-hoc. *, P < 0.05; **, P < 0.01; ****, P < 0.0001.

Nox4 protein levels were not increased in Nox4(f/f) mice at 56 days after SCI, in contrast to our prior observations in spinal cord transected rats [21]. Mean Nox4 protein levels were reduced in muscles from Nox4 cKO mice (Fig. 4A) although some variability was observed. Co-immunoprecipitation experiments were performed to determine if binding of calstabin 1 to RyR1 was altered after SCI in Nox4 cKO mice as compared to Nox4(f/f). There was no difference observed in binding of calstabin 1 to RyR1 when comparing Nox4(f/f) sham and SCI groups (Fig. 4B). Calstabin 1 binding was not significantly different in SCI Nox4 cKO mice compared to sham Nox4 cKO mice (Fig. 4C) likely due to Calstabin 1 interactions with other cell types in skeletal muscle.

**Figure 4.**
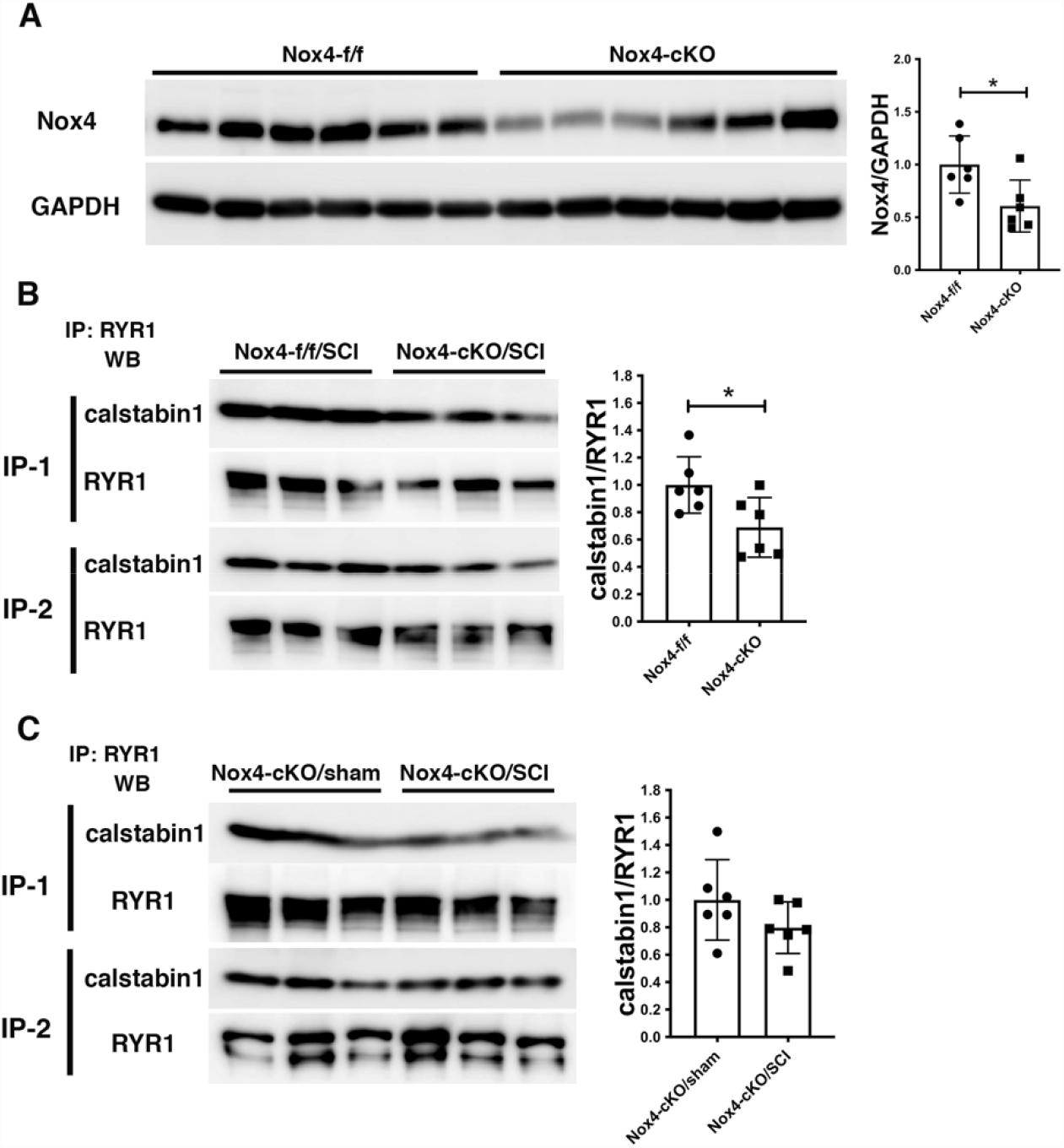
Analysis of Nox4 expression and binding of calstabin 1 to RyR1. A) Nox4 protein levels were determined in whole lysates of gastrocnemius muscle from Nox4 (f/f) and Nox4 cKO by Western blotting s (n=6/group); the graphs show relative abundance of Nox4 after normalization to GAPDH. B) RyR1 was immunoprecipitated from homogenates of gastrocnemius muscle from Nox4(f/f) and Nox4 cKO mice 56 days after SCI (N=6/group) and the immunoprecipitated material analyzed by western blotting for calstabin 1 and RyR1. The graph shows the amount calstabin 1 normalized to the amount of immunoprecipitated RyR1. C) RyR1 was immunoprecipitated from gastrocnemius muscle collected at 56 days after SCI from Nox4 cKO mice (N=6/group) and the immunoprecipitated material subjected to Western blotting for calstabin 1 and RyR1; the graph shows the amount of calstabin 1 normalized to the immunoprecipitated RyR1. Differences between means were evaluated using a two tailed unpaired t-test. *, P < 0.05.

Protein carbonyls are produced by oxidation of amino acids such as lysine, arginine, proline, and threonine. Oxidation by ROS, or Nox4, results in reactive ketones and aldehydes. Protein carbonyls were quantified in the TA muscle of SCI Nox4 cKO, SCI Nox4(f/f), sham Nox4 cKO, and sham Nox4(f/f) mice. Nox4(f/f) TA had significantly more carbonylated protein than Nox4 cKO mice, regardless of whether the animals received SCI or not (F=107.65, p<0.0001) (Fig. 5). Nox4-expressing mice were sensitive to protein oxidation as compared to mice with a cKO of Nox4, suggesting that the oxidase is a significant source of ROS and contributes to muscle weakness.

**Figure 5.**
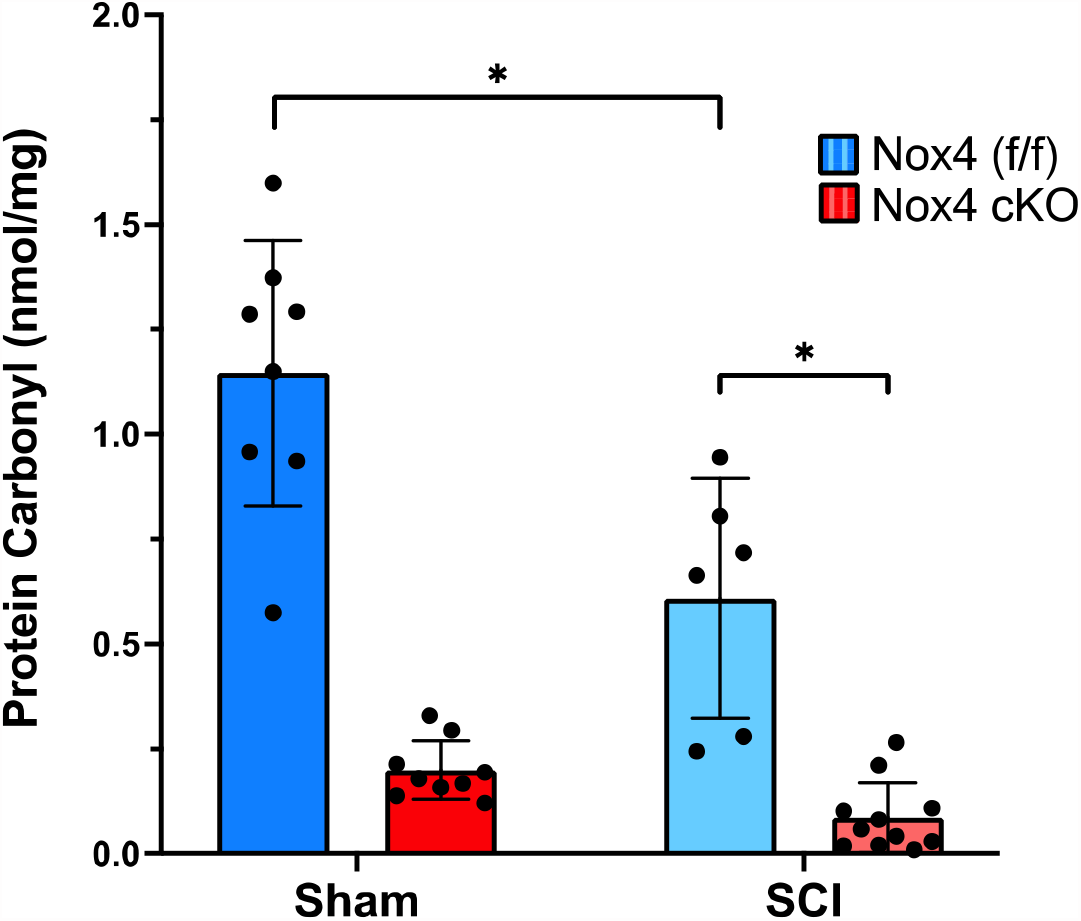
Nox4 increases oxidation of skeletal muscle proteins. TA muscle from controls (Nox4(f/f)) and Nox4 cKO were subjected to either sham surgeries or transection SCI. Protein carbonylation, a result of oxidation, was quantified by ELISA. Nox4(f/f) animals produce significantly more protein carbonyls than Nox4 cKO animals, whether they received sham or SCI surgeries. Dark and light blue bars indicate Nox4 (f/f) Sham and SCI, respectively. Red and pink bars indicate Nox4 cKO Sham and SCI, respectively. Data were analyzed by mixed-model 2-way ANOVA with a Sidak’s multiple comparisons test post-hoc. *, P < 0.05.

Sham Nox4(f/f) mice expressed significantly more carbonylated proteins in TA muscles than SCI Nox4(f/f) mice, indicating that Nox4 produces a considerable amount of ROS even under normal conditions (p<0.0002). Likewise, sham Nox4 cKO mice appear to express more carbonylated proteins than SCI Nox4 cKO mice, but this difference did not reach the level of significance (p=0.6).

## DISCUSSION

The major conclusion supported by the data is that conditional inactivation of Nox4 in skeletal muscle improved muscle force generation in hindlimb muscles from mice with spinal cord transection at 56 days after the SCI, and Nox4 generates some manner of oxidative stress in both healthy sham and SCI mice. The data do not however support mechanistic linkages between Nox4 levels, RyR1-calstabin 1 binding interactions and loss of muscle specific force after SCI. Mechanisms by which reducing Nox4 expression in skeletal muscle increase muscle force generating capacity after SCI cannot be ascertained from the data here presented.

Why our results in Nox4(f/f) mice differ from those in Sprague Dawley rats we previously reported previously [21] is unclear but could reflect species or strain difference in expression levels of regulatory factors that govern Nox4 activity or splice variants that determine how Nox4 is distributed. Nox4 is localized to many cellular compartments that include cytoplasmic membrane, nucleus, mitochondria and endoplasmic reticulum [26]; it remains unclear whether there are species, tissue, or strain dependent differences in Nox4 localization. Distribution of Nox4 in skeletal muscle has been reported [27] but the effects of SCI on Nox4 distribution are yet unknown. While we have found Nox4 bound to RyR1 in lysates of rat skeletal muscle [21], the cell type in which this colocalization occurs remains unknown although we have assumed it to be skeletal muscle fibers. Influence of species, strain or muscle type on expression of Poldip2 or other regulatory factors that govern Nox4 activity is unknown. Nox4 has also been found to be regulated by post-transcriptional modifications such as phosphorylation at Tyrosine 566 by tyrosine kinase Fyn [28]. Effects of SCI on expression or activity of FYN have not been examined. Further studies are needed to address these important questions.

Not much is known about functions of Nox4 in skeletal muscle, and we are aware of only one prior study in which Nox4 was conditionally knocked out in skeletal muscle [27]. The mouse model used in these studies provides a tool for further investigating role(s) of this enzyme in muscle biology. The Nox4 cKO reduced though did not eliminate Nox4 in whole lysates of skeletal muscle which we interpret as indicating expression in cell types within skeletal muscle other than muscle fibers, consistent with the fairly ubiquitous expression of Nox4 [26]. While not the primary focus of our study, it is worth noting that the Nox4 cKO did not significantly alter muscle size or physiological properties in sham-SCI mice although a trend toward a decrease in force production was suggested by the data.

We observed that Nox4(f/f) TA had significantly more carbonylated protein than Nox4 cKO mice, suggesting that the oxidase is a significant source of ROS and contributes to muscle weakness. Sham Nox4(f/f) mice had significantly more carbonylated proteins in TA than SCI Nox4(f/f) mice, indicating that Nox4 produces a considerable amount of ROS even under normal conditions. Likewise, sham Nox4 cKO mice appear to express more carbonylated proteins than SCI Nox4 cKO mice. Two factors may contribute to these findings: first, Nox4 cKO mice express relatively few protein carbonyls overall, resulting in little difference between the sham and SCI animals. Second, SCI results in muscle atrophy, reducing the total amount of muscle proteins, such as RyR1, and increasing the proportion of non-muscle proteins in the lysate [29]. We do not conclude that SCI is protective against Nox4 oxidation, but rather that less Nox4 and fewer proteins available for oxidation are present.

The current study was one of a growing number that have tested whether the skeletal muscle atrophy or reduced skeletal muscle force generating capacity observed after SCI can be blunted by interventions including androgens [30], ActIIB receptor-Fc traps that inactivate myostatin [31], and ursolic acid [32]. Of note, Nox4 cKO is the only intervention shown to increase any measure of specific force generation independently of neurological function. The observation that reduction of Nox4 expression increases muscle force production in completely paralyzed mice offers encouraging evidence that at least some of the defects in skeletal muscle after SCI can be prevented over the long-term. Further study is needed to understand the molecular basis for improved contractile capacity of muscle after SCI in Nox4 cKO mice and whether common mechanisms are operative across species.

## ACKNOWLEDGEMENTS

This work was supported by the Department of Veterans Affairs Office of Research and Development, Rehabilitation Research and Development (RR&D) Service Merit IRX002313-R to CPC, CDA-2 IK2RX002781-01 to ZAG, Center I50RX002020 and by resources provided by the James J. Peters VA Medical Center. NIH grant R01GM 137056 to RI. The work reported herein does not represent the views of the US Department of Veterans Affairs or the US Government.

